# Single-cell atlas of human developing and azoospermia patients’ testicles reveals the roadmap and defects in somatic microenvironment

**DOI:** 10.1101/2020.05.07.082024

**Authors:** LiangYu Zhao, ChenCheng Yao, XiaoYu Xing, Tao Jing, Peng Li, ZiJue Zhu, Chao Yang, Jing Zhai, RuHui Tian, HuiXing Chen, JiaQiang Luo, NaChuan Liu, ZhiWen Deng, XiaoHan Lin, Na Li, Jing Fang, Jie Sun, ChenChen Wang, Zhi Zhou, Zheng Li

**Author notes:** These authors contributed equally (LiangYu Zhao, ChenCheng Yao, XiaoYu Xing, Tao Jing). Correspondence (Z.L.), (Z.Z.), (C.W.), (J.S.).

## Abstract

Non-obstructive azoospermia (NOA) affects 1% of men. However, the unknowns of NOA pathogenesis and even normal spermatogenic microenvironment establishment severely limit the clinical efficacy of NOA treatment. We profiled > 80,000 human testicular single-cell transcriptomes from 10 healthy donors spanning the range from infant to adult and 7 NOA patients. Sertoli cells, which form the scaffold in the testicular microenvironment, exhibited the most obvious damages in NOA patients. We identified the roadmap of Sertoli cell maturation. Notably, Sertoli cells of patients with congenital causes (Klinefelter syndrome and Y chromosome microdeletions) are mature but with abnormal immune response, while the cells in idiopathic NOA (iNOA) are basically physiologically immature. Furthermore, inhibition of Wnt signaling promotes the maturation of Sertoli cells from iNOA patients, allowing these cells to regain their ability to support germ cell survival. We provide a novel perspective on the development of diagnostic methods and therapeutic targets for NOA.

## Introduction

Loss of fertility can be devastating to a patient; unfortunately, 15% of couples in the world are suffering from infertility (Fakhro et al., 2018). In contrast to female infertility, which can often be treated by hormone therapy to stimulate oocyte production, male infertility caused by spermatogenesis abnormalities are much more difficult to treat (Tournaye et al., 2017). Non-obstructive azoospermia (NOA) is the most serious form of male factor infertility, occurring in 10%–15% of infertile men (Dabaja and Schlegel, 2012; Fakhro et al., 2018). However, only a small percentage is caused by congenital factors such as Klinefelter syndrome (KS) and Y chromosome AZF region microdeletion (AZF_Del). Most of the remaining cases are due to unknown causes, also known as idiopathic NOA (iNOA), accounting for over 70% (Dabaja and Schlegel, 2012). Although the final morphological feature in iNOA testis is that there are no or few spermatogenic cells, sperm can be found in 10% of iNOA patients by surgery (Dabaja and Schlegel, 2012). Notably, the degree of testicular development may predict adverse patient outcomes. Hence, effective etiology analysis and therapies for patients with NOA are urgently required.

Spermatogenesis depends on the full maturation of the somatic microenvironment. It is an extremely complicated process that occurs after birth and is completed after puberty. An altered microenvironment, which might affect fertility, has been observed in NOA patients (Ma et al., 2013; McLachlan et al., 2007). However, our understanding of cells in the somatic microenvironment remains limited. We have a superficial understanding that Sertoli cell development after birth is mainly divided into two stages: the immature and mature stage. Maturation is regulated by the dramatic changes in hormone levels during puberty (Simorangkir et al., 2012). Little is known about this maturation process, Sertoli cell heterogeneity, and interaction with other cell types at the molecular level. The same is true for other testicular cell populations, hampering our understanding of the pathogenesis of spermatogenic disorders.

The rapid development of single-cell RNA sequencing allows us to investigate individual cell populations in testis at unprecedented resolution. Most previous studies have focused on germ cell spermatogenesis itself (Chen et al., 2018; Wang et al., 2018). A recent study of human testicular single-cell transcriptomes in puberty has revealed developmental changes in somatic cells (Guo et al., 2020). However, we still do not know the major differentiation signals, metabolic characteristics, and cell–cell interactions of these cells at various developmental stages. Here, we profiled 88,723 individual testicular cells from 10 healthy subjects of various ages and 7 patients with one of the three most common types of NOA. Focusing on the somatic cell dataset, we identified three independent stages during Sertoli cell maturation and uncovered the pathological changes and maturation disorders in different types of NOA Sertoli cells. In addition, we obtained evidence that the Wnt signaling pathway regulates the maturation of Sertoli cells in both normal and NOA patients. Collectively, our results provide in-depth insight into the maturation of the spermatogenic microenvironment and the mechanisms underlying pathogenesis, thereby offering new targets for NOA treatment strategies.

## Results

### Overview of the hierarchies of multiple cell populations in healthy and NOA human testes

To characterize the baseline cellular diversity of testicles during human developing and under pathological state, we profiled cells from 10 donors with normal spermatogenesis (aged 2–31 years) and 7 NOA patients (KS (3 cases), AZFa_Del (1 case), and iNOA (3 cases)) (Figure 1A and S1A). The sex hormone levels of 10 physiological donors were within the normal range for their respective ages (Condorelli et al., 2018) (Figure S1B). The PAS/hematoxylin staining results showed that all 10 physiological testes had a normal morphology with their ages (Figure S1C). In addition, the spermatogenic stages were identified by germ cell markers, including GFRA1 (spermatogonial stem cells), c-KIT (differentiating spermatogonia), SYCP3/γH2AX (spermatocytes), and PNA (spermatids/spermatozoa). These spermatogenic stages reflected the ages of the healthy donors (Figure S1D). Uniform manifold approximation and projection (UMAP) analysis of these cells showed that the heterogeneity in different age groups was obvious, but all 5 healthy adult samples had a good repeatability (Figures 1B and S2D). On average, 9359 reads (UMIs), 2719 expressed genes and 3.7% mitochondrial genes were detected per cell (Figure S2A).

**Figure 1.**
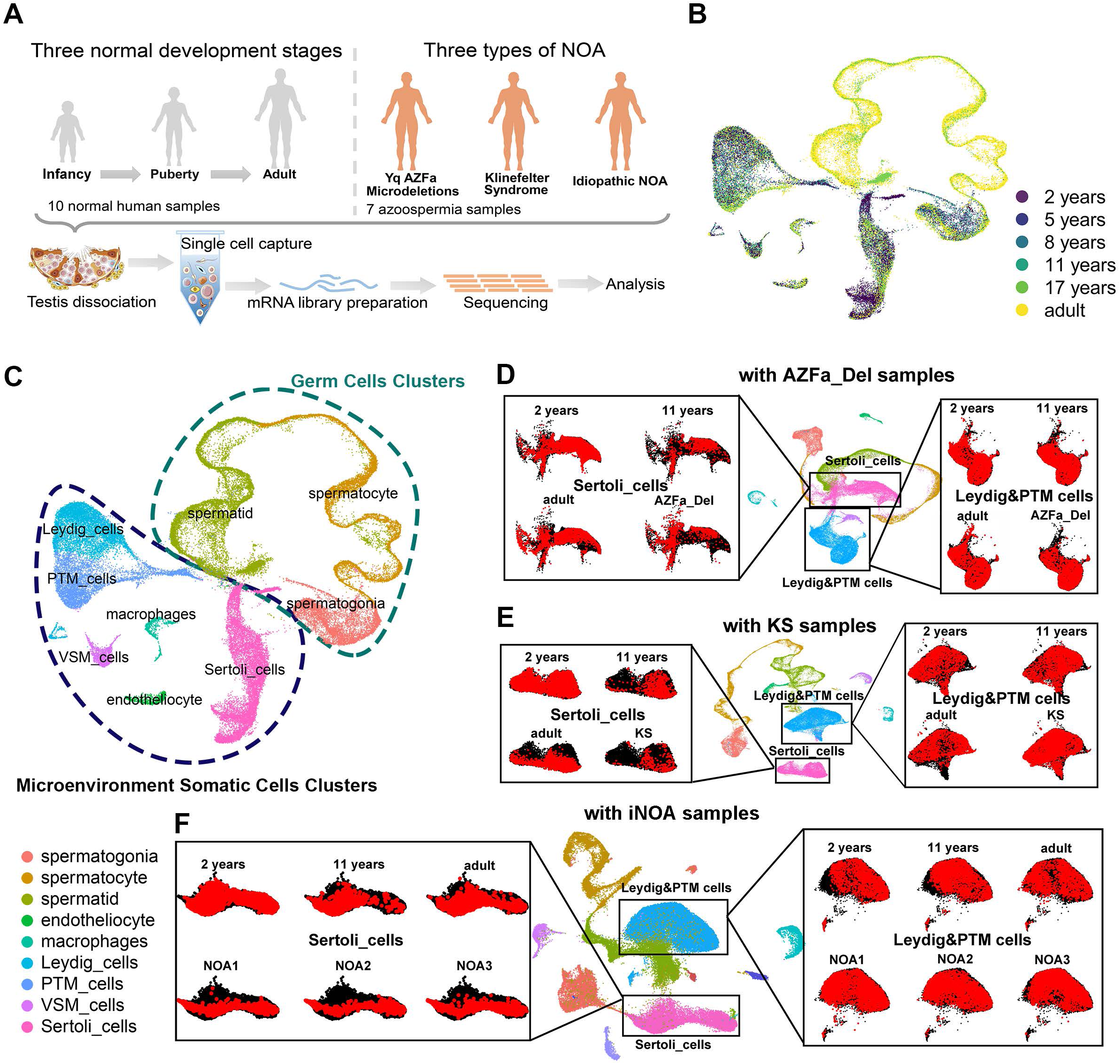
Global expression profiling of human testicular cells from infancy to adulthood and in NOA patients by single-cell RNA-seq. (A) Schematic illustration of the experimental workflow. (B, C) UMAP plots of all testicular cells from 10 healthy subjects. Cells are colored for (B) ages or (C) types. UMAP, uniform manifold approximation and projection. (D–F) UMAP plots of all testicular cells from 10 healthy subjects merged with (D) 1 AZFa_Del sample, (E) 3 KS samples, or (F) 3 iNOA samples. Sertoli cells (left panels) or Leydig&PTM cells (right panels) were isolated and are highlighted as red according to the sample type. PTM cell, peritubular myoid cell; VSM cell, vascular smooth muscle cell.

A total of nine cell clusters were identified in the whole cell population of 10 healthy subjects based on the expression of known cell type-specific markers (Figure S2G). These nine clusters could be further divided into two parts, germ cells and microenvironment somatic cells (Figure 1C). The latter included endotheliocytes, macrophages, vascular smooth muscle cells (VSM_cells), peritubular myoid (PTM) cells, and Leydig cells. Total 1101 differentially expressed genes (DEGs) with a fold change of log2 transformed UMI > 1 of each cluster were identified. The results of our gene ontology (GO) analysis of these DEGs were consistent with our understanding of the biological processes of these cell types (Table 1, Figure S2E-F).

To gain insight into the cellular differences between testes form healthy subjects and NOA patients, we re-clustered testicular somatic cells from 10 healthy donors with AZFa_Del, KS, and iNOA (Figure 1D–F). Sertoli cells showed a greater difference in spatial distribution than other cell populations on UMI plot (Figure 1D–F). Considering the fact that Sertoli cells, which are located around and are in direct contact with germ cells, function as scaffolds and “nurse” cells in the spermatogenic microenvironment (Hai et al., 2014), our results suggest that the dysfunction of Sertoli cells contributes to the failure of the establishment of a suitable microenvironment in NOA.

### Identification of three stages and heterogeneity of human Sertoli cells during maturation

To explore the heterogeneity of Sertoli cells during normal development, we re-clustered the Sertoli cells and identified three subpopulations in 10 healthy subjects (Figure 2A-B). To analyze the origin and maturation process of Sertoli cells, we performed pseudotime trajectory analysis based on clustering combined with the label of “age.” (Figure 2A). Most Sertoli cells in 2-year-olds were identified as Stage_a cells, and the ratio between the numbers of Sertoli cells in Stage_a and Stage_b declined with age and reached the lowest point at puberty (11 years). Stage_c Sertoli cells did not appear until after the age of 11 years and were predominant in the late puberty and adult. It is worth noting that the full range of Sertoli cell stages was observed in adult testes (Figure 2C), suggesting that the stepwise maturation of Sertoli cells starts at a very young age, and reaches a steady state in late puberty.

**Figure 2.**
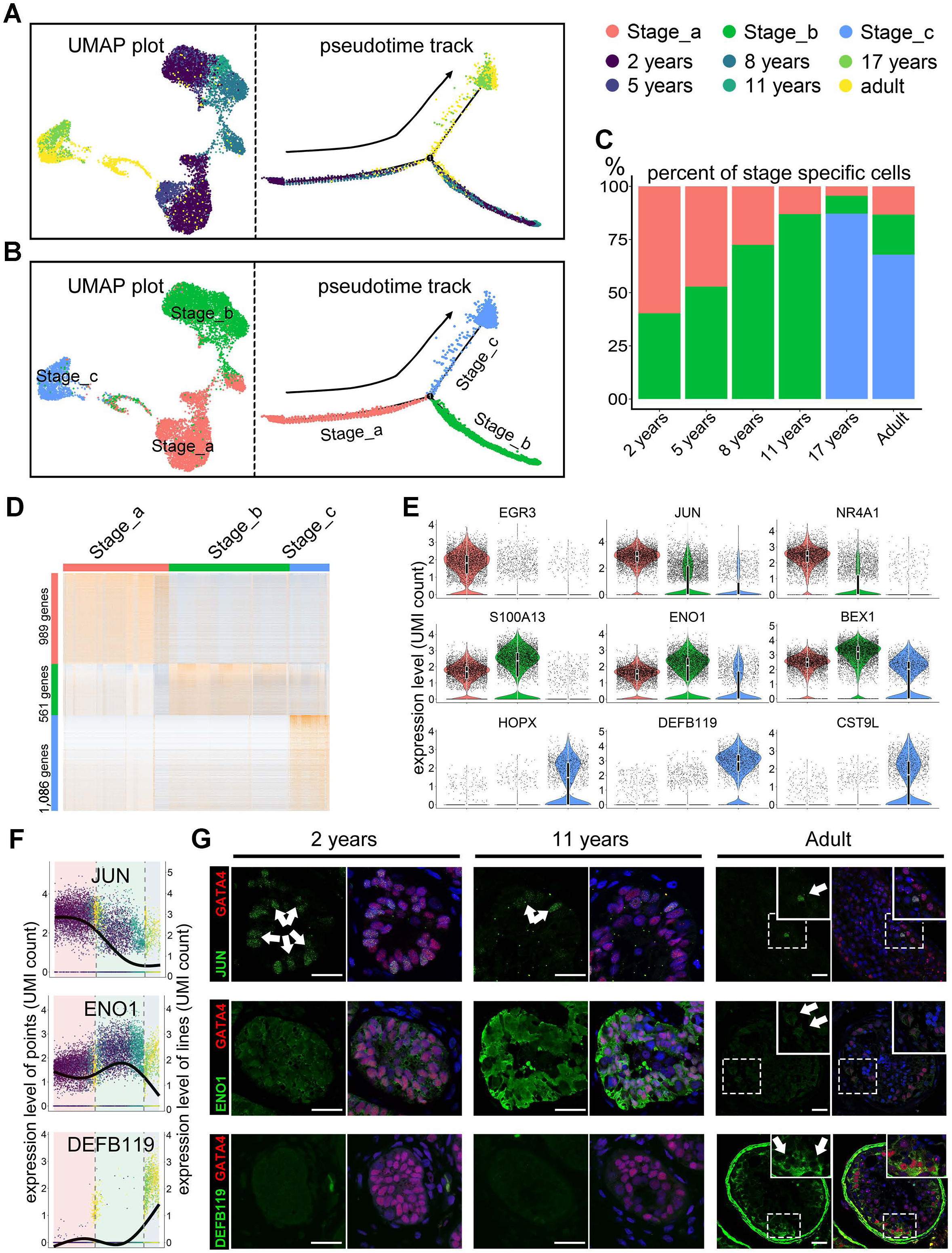
Identification of three maturation stages during Sertoli cell development. (A, B) Analysis of Sertoli cells (UMAP plot and pseudotime trajectory plot), with cells colored by (A) age or (B) stage. (C) Bar plot showing the proportion of Sertoli cells at each stage (Stage_a, red; Stage_b, green; Stage_c, blue) in each age group. (D) Heatmap showing the DEGs of each stage during Sertoli cell maturation. DEG counts are shown on the left of the color bar of the cell type annotation. (E) Violin plot showing the expression levels of the top DEGs at each stage (Stage_a, red; Stage_b, green; Stage_c, blue). (F) Expression levels of *JUN, ENO1*, and *DEFB119* in Sertoli cells ordered in pseudotime. (G) Immunofluorescence co-staining of GATA4 (green) with JUN (red, upper panel), ENO1 (red, middle panel), and DEFB119 (red, lower panel) in human testicular paraffin sections at three ages. The scale bar represents 20 μm.

Next, we further analyzed the dynamic changes in the gene expression pattern at each stage. The total number of expressed genes and total UMI count per cell declined dramatically from Stage_a to Stage_b and Stage_c (Figure S3A). 989, 561, and 1086 DEGs were observed in three states, respectively (Table S2 and Figure 2D). *EGR3, JUN*, and *NR4A1* were the top three DEGs in Stage_a, *S100A13, ENO1*, and *BEX1* were the top three DEGs in Stage_b, and *HOPX, DEFB119*, and *CST9L* were the top three DEGs in Stage_c (Figure 2E). To verify this result, we performed immunohistochemical staining of testis sections from subjects of different ages for JUN, ENO1, and DEFB119 (Figure 2F-G). Then, we performed GO analysis of DEGs at each stage. In Stage_a cells, the main enriched GO terms were “stem cell differentiation,” “cell fate commitment,” and “maintenance of cell number” (Figure S3B), suggesting that at this stage Sertoli cells exhibit some characteristics of stem or progenitor cells. In Stage_b, “small molecule catabolic process,” “generation of precursor metabolites and energy,” and “cellular amino acid metabolic process” were enriched (Figure S3C). In Stage_c, the main enriched GO terms were “cellular metabolic compound salvage,” “protein transmembrane transport,” and “phagosome maturation” (Figure S3D), indicating that the main functions of mature Sertoli cells are phagocytosis of germ cells and their metabolite.

Then, we proceeded to annotate basic physiological characteristics, and compared the cell cycle state and the energy metabolism type of the three stages (Figure S3E-G). Based on the previously reported S and G2/M phase-specific genes (Nestorowa et al., 2016), we found Sertoli cells in Stage_a expressed higher levels of mitotic genes than other two later stages (Figure S3E). After calculating the mitotic phase score for each cell based on Seurat R package, we estimated that Stage_a contain the highest proportion of S phase cells (Figure S3F). As regards heterogeneity in energy metabolism, the expression levels of glycolysis-related genes increased from Stage_a to Stage_c; however, the oxidative phosphorylation-related genes showed an opposite trend. In addition, the expression levels of triglyceride metabolism-related genes in Sertoli cells were higher than in the other eight types of testicular cells, and it was also higher in Stage_b and Stage_c than in Stage_a (Figure S3G).

All these changes in expression profile, proliferation and energy metabolism revealed that Sertoli cells went through three distinct consecutive developmental stages.

### Regulatory networks during Sertoli cell maturation

We hypothesized that the levels of key regulators should change dramatically at the junction between two consecutive stages. Therefore, we screened 372 candidate regulators that showed a significant change at a branch point of the pseudotime trajectory (Figure 3A). GO terms associated with these genes included “TGF-beta signal pathway,” “tube development,” and “response to steroid hormone” (Figure 3B), indicating steroid hormones are one of the main upstream extracellular regulatory signals. The heatmap of “response to steroid hormone”-related genes is shown in (Figure 3C). In addition, we identified pathways that potentially regulate the maturation of Sertoli cells by Ingenuity Pathway Analysis (IPA). IPA analysis showed that from Stage_a to Stage_b, “ERK5 signaling,” “IGF-1 signaling,” and “EGF signaling”, et al were inhibited, while “remodeling of epithelial adherens junctions”, et al were activated (Figure 3D). In addition, from Stage_b to Stage_c, “cardiac hypertrophy signaling,” and “Wnt/β-catenin signaling” et al were inhibited and “germ cell–Sertoli cell junction signaling” et al were activated (Figure 3E). These results indicate that the proliferation of Sertoli cells mainly occurs in Stage_a, and structural remodeling mainly occurs in Stage_b.

**Figure 3.**
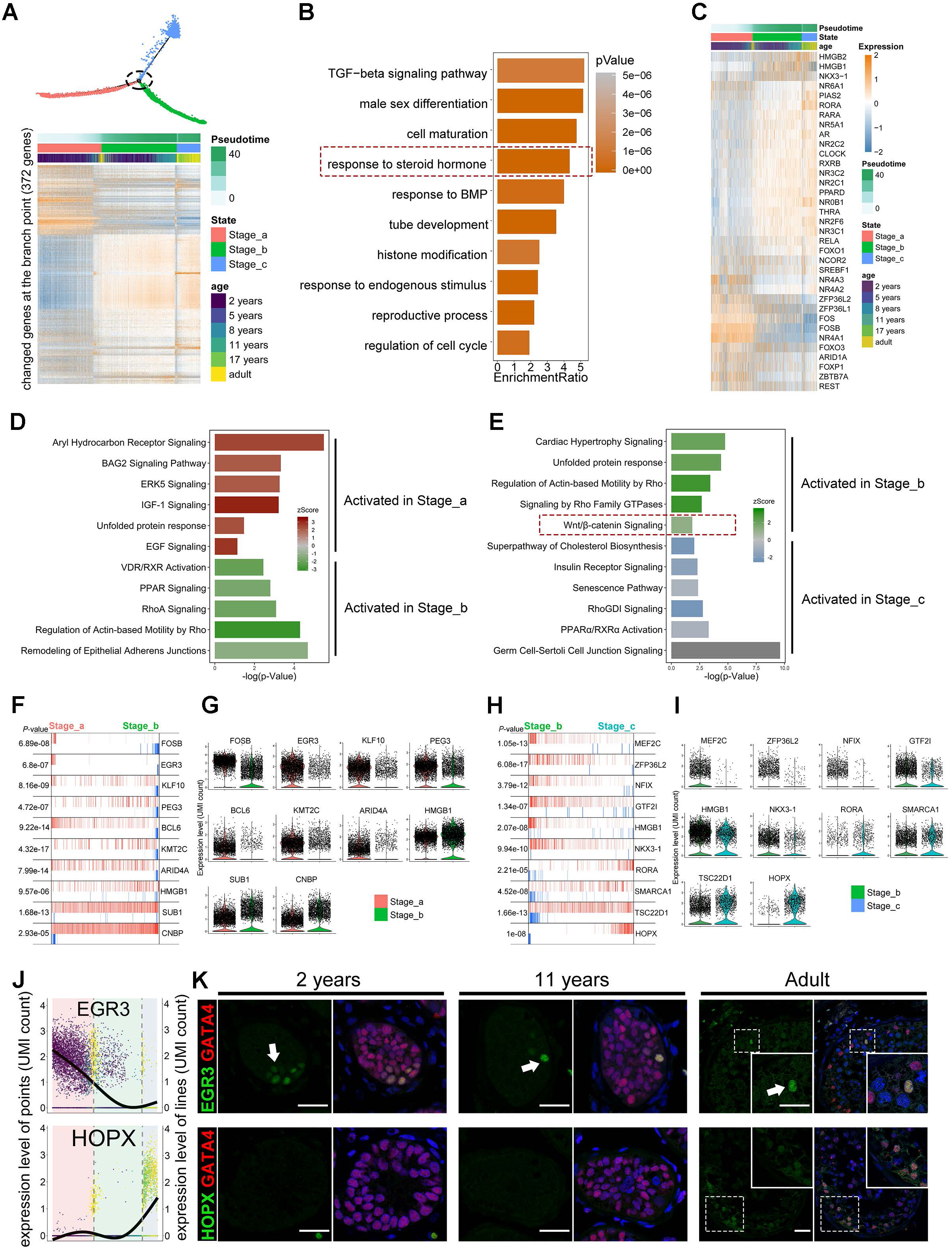
Inferred signal pathways and master regulators for three stages of Sertoli cells. (A) Heatmap of 372 genes that exhibited dramatic changes in gene expression at the intersection of two consecutive stages of Sertoli cell development. (B) Pathways and biological process terms of GSEA are shown as barplot. The GSEA score is presented on the *x*-axis, and the gradient of orange indicates low to high FDR values. (C) Heatmap of 37 genes related to the GO term “response to steroid hormone.” (D, E) Pathway terms of IPA enriched at each stage are shown as barplot. The *P*-value is presented on the *x*-axis, and the gradient of two colors indicates low to high activation scores in the two stages. (F-H) The top 10 candidate master regulators at each stage of Sertoli cell development identified by the master regulator analysis algorithm (MARINa) and (G, I) violin plots of the expression levels of those regulators in two consecutive stages. In the MARINa plots, activated targets are colored red and repressed targets are colored blue for each potential master regulator (vertical lines on the *x*-axis). On the *x*-axis, genes were rank-sorted by their differential expression in two consecutive developmental stages. The *P*-values on the left indicate the significance of enrichment, calculated by permutating two developmental phases. (J) Expression levels of *EGR3* and *HOPX* in Sertoli cells ordered in pseudotime. (K) Immunofluorescence co-staining of GATA4 (red) with a master regulator of Stage_a, EGR3 (green, upper panel), and a master regulator of Stage_c, HOPX (green, lower panel), in human testicular paraffin sections at three ages.

To explore the key regulators and identify the transcriptional regulatory network during Sertoli cell maturation, we analyzed 1665 human transcription factors using the Algorithm for the Reconstruction of Accurate Cellular Networks (ARACNe) (Lachmann et al., 2016), and the *ssmarina* package in R was used to analyze their downstream gene set. We found that FOSB, EGR3, KLF10, et al might specifically play a critical role in Stage_a Sertoli cells (Figure 3F-G), whereas HMGB1, SUB1, CNBP,et al might be the master regulators in Stage_b Sertoli cells (Figure 3F–I). In Sertoli cells in Stage_c, RORA, SMARCA1 and HOPX were the top candidate gene expression regulators (Figure 3H-I). These results were confirmed by staining testis sections for EGR3 and HOPX. On average, there were 2.35 EGR3-positive Sertoli cells per tubule in testis sections of 2-year-olds, which was significantly higher than in 11-year-olds (0.54) and adults (0.05) (Figure 3K). No HOPX-positive Sertoli cells were detected in testis sections from 2- and 11-year-olds, whereas in sections from 17-year-olds and adults, nearly all Sertoli cells were HOPX-positive (Figure 3K). These results provide novel insight in potential regulatory networks, ranging from the upstream hormonal signals to intracellular regulatory pathways.

### The multi-lineage interactome network and its dynamic changes in the spermatogenic microenvironment

To investigate the complex signaling networks and their dynamic changes in the spermatogenic microenvironment, we performed an unbiased ligand–receptor interaction analysis between these testicular cell subsets by CellphoneDB (Vento-Tormo et al., 2018). In Stage_a, Stage_b, and Stage_c, we found 158, 96, and 72 interactions between Sertoli cell ligands and receptors from other cells and 189, 49, and 41 interactions between ligands from other cells and Sertoli cell receptors, respectively (Figure S4A–C). The interaction between Sertoli cells and other testicular cells showed unique changes in different stage and cell type (Figure S4D). Focusing on the interaction between Sertoli cells and spermatogonia, we found most interaction pairs involved in TGF-β signaling. The expression of the ligands of TGF-β signaling decreased rapidly after puberty (Figure S4E). Interestingly, in most interaction pairs between germ cells and Sertoli cells, Sertoli cells play the role of signal source, whereas in most interaction pairs between LEYDIG&PTM cells and Sertoli cells, Sertoli cells receive signals in the early stage. IGF and NOTCH signaling between Sertoli cells and LEYDIG&PTM cells play important roles in the early stage of spermatogenic microenvironment maturation (Figure S4D). Interactions between collagen (COL1A2) in Sertoli cells and integrins α1β1, α10β1, and α2β1 in endotheliocytes were detected at the early stage. In addition, Sertoli cells also expressed the ligands of CD74, which is mainly expressed in macrophages; this interaction occurs mainly in Stage_b (Figure S4D)

These findings indicate complex paracrine regulatory networks of the maturation of the human spermatogenic microenvironment.

### Single-cell profiling revealed that different types of NOA testes have distinct features of Sertoli cell defects

An altered microenvironment has been reported in NOA patients in previous studies (Ma et al., 2013; Tarulli et al., 2014), but the underlying pathological mechanisms remain unknown. Therefore, we re-clustered 10 normal samples with samples from 3 iNOA patients, 3 KS patients, and 1 AZFa_Del patient to investigate the changes of the microenvironment under pathologic states (Figure S5A-B). Interestingly, we found that samples from different pathological types showed significant differences, whereas low heterogeneity was observed among samples belonging to the same type (Figure S5C). In addition, the quality of single-cell sequencing libraries (total UMI count and number of expressed genes per cell) in all three pathological types was within an acceptable range (Figure S5D).

Focusing on Sertoli cells, we found that part of the iNOA Sertoli cells overlapped with normal immature Sertoli cells; however, Sertoli cells of AZFa_Del and KS patients were close to but did not overlap with any normal immature samples in the UMAP reduced-dimension plot (Figure 4A-B). Pseudotime trajectory analysis showed that iNOA Sertoli cells were considerably heterogeneous, as one cluster was in Stage_a and another in Stage_b; AZFa_Del Sertoli cells were in the early part of Stage_c; and KS Sertoli cells were arrested in Stage_b (Figure 4B). Cell cycle analysis showed iNOA Sertoli cells contained more S phase cells than other adult objects (Figure 4C), suggesting a higher proliferation rate in iNOA Sertoli cells. In addition, Sertoli cells in the three types of NOA showed energy metabolism patterns that were different from that of healthy adults. The expression levels of glycolysis-related genes in both KS (0.73%) and AZFa_Del (0.64%) Sertoli cells were higher than in iNOA (0.41%) and healthy adult (0.42%). The expression levels of oxidative phosphorylation-related genes in iNOA (1.31%) Sertoli cells were much lower than in healthy adult (5.5%), KS (5.8%), and AZFa_Del (4.9%) Sertoli cells. Triglyceride metabolism is important for Sertoli cells to perform physiological functions. Our data showed that AZFa_Del Sertoli cells (0.68%) exhibit the highest triglyceride metabolic activity, followed by healthy adult (0.35%), KS (0.37%), and AZFa_Del (0.25%) Sertoli cells (Figure 4D). Some important regulators of Sertoli cell development were also shown to be differently expressed between NOA patients and healthy subjects; for example, expression of the Stage_a regulator gene *FOSB* was higher in iNOA and AZFa_Del, and *EGR3* was highly expressed in iNOA, while the transcriptional activity of the Stage_c regulators *HOPX* and *TSC22D1* in iNOA was lower than in other adult samples (Figure 4E). This result was corroborated by immunofluorescence co-staining of HOPX with the Sertoli cell marker GATA4; HOPX levels were lower in iNOA samples than in samples from other patients and healthy adults (Figure 4F).

**Figure 4.**
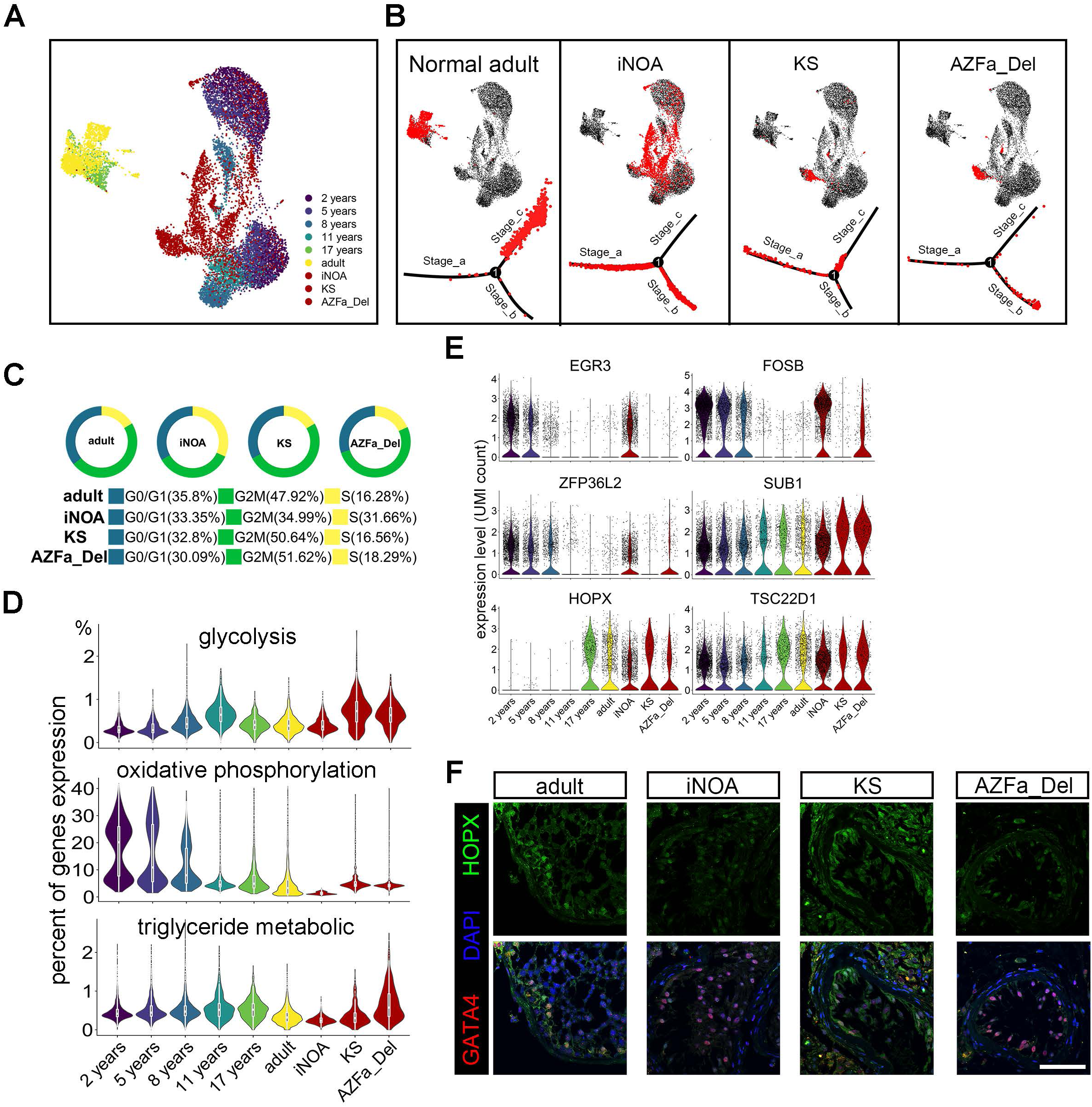
Single-cell transcriptome profiling of three types of NOA Sertoli cells. (A) Analysis of normal Sertoli cells at six ages merged with iNOA, AZFa_Del, and KS Sertoli cells, in a UMAP plot with cells colored by age and sample type. (B) Healthy adult, iNOA, AZFa_Del, and KS Sertoli cells are highlighted red in the UMAP plot (upper row) and the pseudotime trajectory plot (lower row). (C) Cell cycle score of healthy adults and three types of pathological Sertoli cells. (D) Violin plot of the expression levels of energy metabolism-related genes in healthy (six different ages) and three types of NOA Sertoli cells. (E) Violin plot of the expression levels of stage-specific master regulators in healthy (six different ages) and three types of NOA Sertoli cells. (F) Immunofluorescence co-staining of GATA4 (red) and HOPX (green) in healthy adult and three types of pathological testicular paraffin sections. The scale bar represents 50 μm.

### Sertoli cells of KS and AZFa_Del patients exhibited abnormal gene expression patterns

KS and AZFa_Del patients have clear etiologies (disorder of the sex chromosomes at the gene or chromosome level), but unknown pathogenesis for spermatogenic dysfunction. Therefore, we analyzed the differences in sex chromosome gene expression patterns of these two NOA types and compared them with those of healthy subjects to analyze potential mechanisms of pathogenesis. PAS staining showed that the AZFa_Del sample was Sertoli cell-only syndrome (SCOS) with a Johnsen score of 2 (Figure S1C). In total, 1286 and 1426 DEGs were down- and upregulated in AZFa_Del Sertoli cells, respectively, compared with control adult Sertoli cells (Table S4 and Figure S6A). Gene set enrichment analysis (GSEA) terms such as “chromosome localization” and “double-strand break repair” were decreased in AZFa_Del, while “humoral immune response” and “regulation of immune system process” were enriched (Figure S6B). In addition, genes such as macrophage migration inhibitory factor (MIF) and defensin beta 119 (DEFB119), involved in cell-mediated immunity, immunoregulation, and inflammation, were also highly expressed in AZFa_Del (Figure S6C), suggesting aberrant immune activation is involved in the pathogenesis of AZFa_Del Sertoli cells. Interestingly, the expression percentage of X chromosomal but not Y chromosomal genes in AZFa_Del Sertoli cells was different from healthy adults (Figure S6D). The AZFa region contains three genes, i.e., *USP9Y, DBY*, and *UTY*, among them, *USP9Y* was highly expressed in Sertoli cells (Figure S6E). All three AZFa genes were highly expressed before puberty and were downregulated with age. Low expression of *UTY* was detected in AZFa_Del Sertoli cells, indicating there are multiple copies of this gene outside the AZFa region (Figure S6F).

In KS testes, most seminiferous tubules suffered more severe atrophy than AZFa_Del, with Johnsen scores of 0–1, and exhibited vascular basement membrane thickening combined with severe interstitial proliferation, while the remaining seminiferous tubules contained few Sertoli cells with abnormal morphology, as shown by testicular biopsy (Figure S1C), indicating the abnormal X chromosomes affected both germ cells and Sertoli cells. Total 831 and 1130 DEGs were down- and upregulated in KS Sertoli cells compared with healthy adults (Table S5 and Figure S6G). Interestingly, immune response-related terms such as “response to interferon-alpha” and “humoral immune response” were positively enriched in KS, similar to AZFa_Del (Figure S6H). In addition, *MIF* and beta-2-microglobulin (*B2M*) were found among the top DEGs in KS (Figure S6I), indicating aberrant immune activation might be a universal pathogenic process resulting from sex chromosome dysfunction. A subset of X-linked genes escapes silencing by X-inactivation and is expressed from both X chromosomes. Therefore, we analyzed escaping X-linked genes (eXi genes) and non-escaping X-linked genes (neXi genes) separately (Carrel and Willard, 2005). The expression of eXi genes in KS was higher than in healthy adults in all eight types of testicular cells (Figure S6J). Higher expression levels of neXi genes were observed in germ cells and Sertoli cells of KS patients, but these differences were not found in endotheliocytes, macrophages, and Leydig&PTM cells (Figure S6J). The fold changes of eXi genes and neXi genes were calculated (Figure S6L). With respect to Sertoli cells, the expression of both eXi genes and neXi genes in KS Sertoli cells was higher than in healthy adults (Figure S6K).

### Sertoli cell profiling revealed signals involved in maturation defects in iNOA patients

The etiology of iNOA could not be explained by genetic abnormalities of sex chromosomes or otherwise. Therefore, we further investigated the iNOA Sertoli cells, focusing on the heterogeneity of maturation, the proliferation ability, and their regulatory network. PAS staining of testicular biopsies of all 3 iNOA patients revealed typical SCOS with Johnsen scores of 2; vascular basement membrane thickening and over proliferation of interstitial cells were also observed, but these were not as severe as in KS (Figure S1C). The UMAP plot of Sertoli cells from 5 healthy adults and 3 iNOA patients showed that these Sertoli cells could be further clustered into three subsets, i.e., clusters 1 and 2 (composed of iNOA Sertoli cells) and cluster 3 (healthy adult Sertoli cells) (Figure 5A). Pseudotime trajectory analysis showed the maturation characteristics of cluster 1 and cluster 2 were close to those of Stage_a and Stage_b Sertoli cells, respectively (Figure 5B). The heterogeneity of iNOA Sertoli cells was confirmed by the heatmap of DEGs between healthy adults and iNOA patients (Figure 5C). Of the DEGs in cluster 1, 52.6% overlapped with those in Stage_a, while only 15.6% overlap was observed between cluster 2 and Stage_b and between cluster 3 and Stage_c (Figure 5D), indicating that cluster 1 contained immature Sertoli cells that resembled Stage_a cells, but cluster 2 contained immature Sertoli cells with a pathological transcription profile. The top DEGs of each stage during Sertoli cell development coincided, at least partially, with the cluster-specific expression patterns of the three clusters (Figure 5E). Next, we performed GSEA using the DEGs of clusters 1 and 2. The GSEA terms which were enriched in normal Stage_a Sertoli cells, such as “maintenance of cell number” and “stem cell differentiation,” were also enriched in cluster 1 iNOA Sertoli cells (Figure 5F), indicating the immaturity of cluster 1 Sertoli cells. The enriched GSEA terms in cluster 2, such as “phagocytosis,” “immune response-regulating signaling pathway,” and “response to steroid hormone,” also indicated cluster 2 shows similarity with Stage_b cells at some level (Figure 5G).

**Figure 5.**
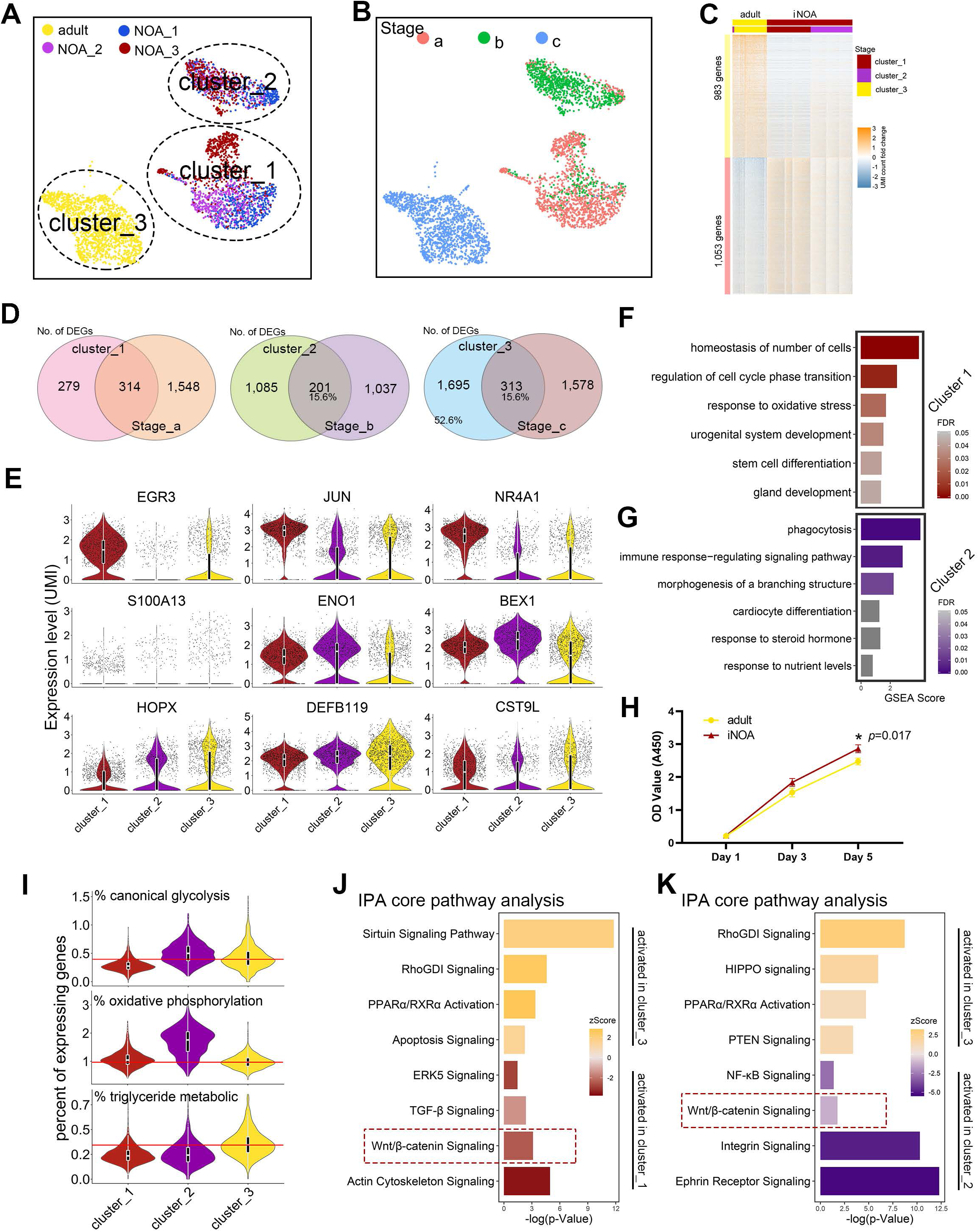
The changes in expression pattern showed the heterogeneity and maturation arrest of iNOA Sertoli cells. (A, B) UMAP plot showing the heterogeneity among iNOA Sertoli cells; a clear difference was observed between iNOA and normal adult Sertoli cells. (C) Heatmap of DEGs between healthy adult and iNOA Sertoli cells. (D) Venn diagram showing the overlap between DEGs in each cluster and in the three developmental stages. (E) Violin plot of the expression levels of candidate stage-specific markers in three Sertoli cell clusters. The upper row shows Stage_a markers; the middle row shows Stage_b markers, and the bottom row shows Stage_c markers. (F, G) GSEA terms enriched in (F) cluster 1 and (G) cluster 2 iNOA Sertoli cells, shown as barplot. The GSEA score is presented on the *x*-axis, and the gradient of red indicates low to high FDR values. (H) iNOA Sertoli cells proliferated faster than normal adult Sertoli cells in vitro, as shown by CCK-8 assay. (I) Violin plot of the expression levels of energy metabolism-related genes in three Sertoli cell clusters. (J, K) Pathways terms of IPA enriched in (J) cluster 1 and (K) cluster 2 compared with normal adult Sertoli cells, shown as barplot. The *P*-value is presented on the *x*-axis, and the gradient of two colors indicates low to high activation scores in the two clusters.

We found that iNOA Sertoli cells were more proliferative (Figure 5H). Regarding the energy metabolism pattern, energy metabolism characteristics of clusters 1, 2, and 3 were similar to those of Stage_a, Stage_b, and Stage_c Sertoli cells, respectively, except that cluster 1 showed lower expression of oxidative phosphorylation-related genes than Stage_a cells (Figure 5I). All these results indicate a maturation arrest in part of the iNOA Sertoli cells.

To better understand the mechanisms underlying maturation arrest and aberrant transcription, we compared cluster 3 with cluster 1 and cluster 2, and then performed core IPA with the DEGs. Pathway terms such as “ERK5 signaling,” “TGF-β signaling,” and “Wnt/β-catenin signaling,” were enriched in cluster 1 (Figure 5J). In addition, pathway terms such as “RhoGDI signaling” and “PPARα/RXRα activation” were decreased in clusters 1 and 2, suggesting the silencing of these two pathways may mediate a pathological transformation in iNOA Sertoli cells. We found that “integrin signaling,” “ephrin receptor signaling,” and “Wnt/β-catenin signaling” were activated in cluster 2 iNOA Sertoli cells (Figure 5K). Considering Wnt/β-catenin signaling was inhibited in healthy Stage_b and Stage_c cells (Figure 3D-E) and iNOA Sertoli cells showed reduced maturation, we hypothesized that the aberrant activation of the Wnt pathway may cause maturation defects in iNOA Sertoli cells.

### Inhibition of the Wnt pathway restored the maturation defects of Sertoli cells in iNOA patients

We found mature Sertoli cells would de-differentiate and “re-enter” into Stage_a when isolated from testicular tissue and cultured in vitro. These changes manifested as follows: (1) mature Sertoli cells were in the resting phase but regained proliferation ability in vitro (Figure 6C); (2) the expression of stage-specific markers, such as HOPX and JUN, more closely resembled that of Stage_a Sertoli cells (Figures 2F and 6A-E); and (3) iNOA Sertoli cells were arrested in a relatively naive state in vivo, but they showed no difference in maturation compared with normal adult cells when cultured in vitro. Importantly, we noticed Wnt/β-catenin signaling was activated in immature and iNOA Sertoli cells (Figures 3E and 5J-K) and was inhibited in normal mature Sertoli cells, suggesting the Wnt pathway may play an important role in the regulation of Sertoli cell maturation. To investigate this hypothesis, we cultured iNOA and normal adult Sertoli cells in vitro and treated them with or without ICG-001 (ICG), an inhibitor of the Wnt signaling pathway (Akcora et al., 2018). After 14 days, the ICG-treated Sertoli cells had a larger cell surface, reduced refraction, and an altered internal texture (Figure 6B). In addition, CCK-8 results showed that ICG treatment strongly decreased the proliferation of both iNOA and normal adult Sertoli cells (Figure 6C).

**Figure 6.**
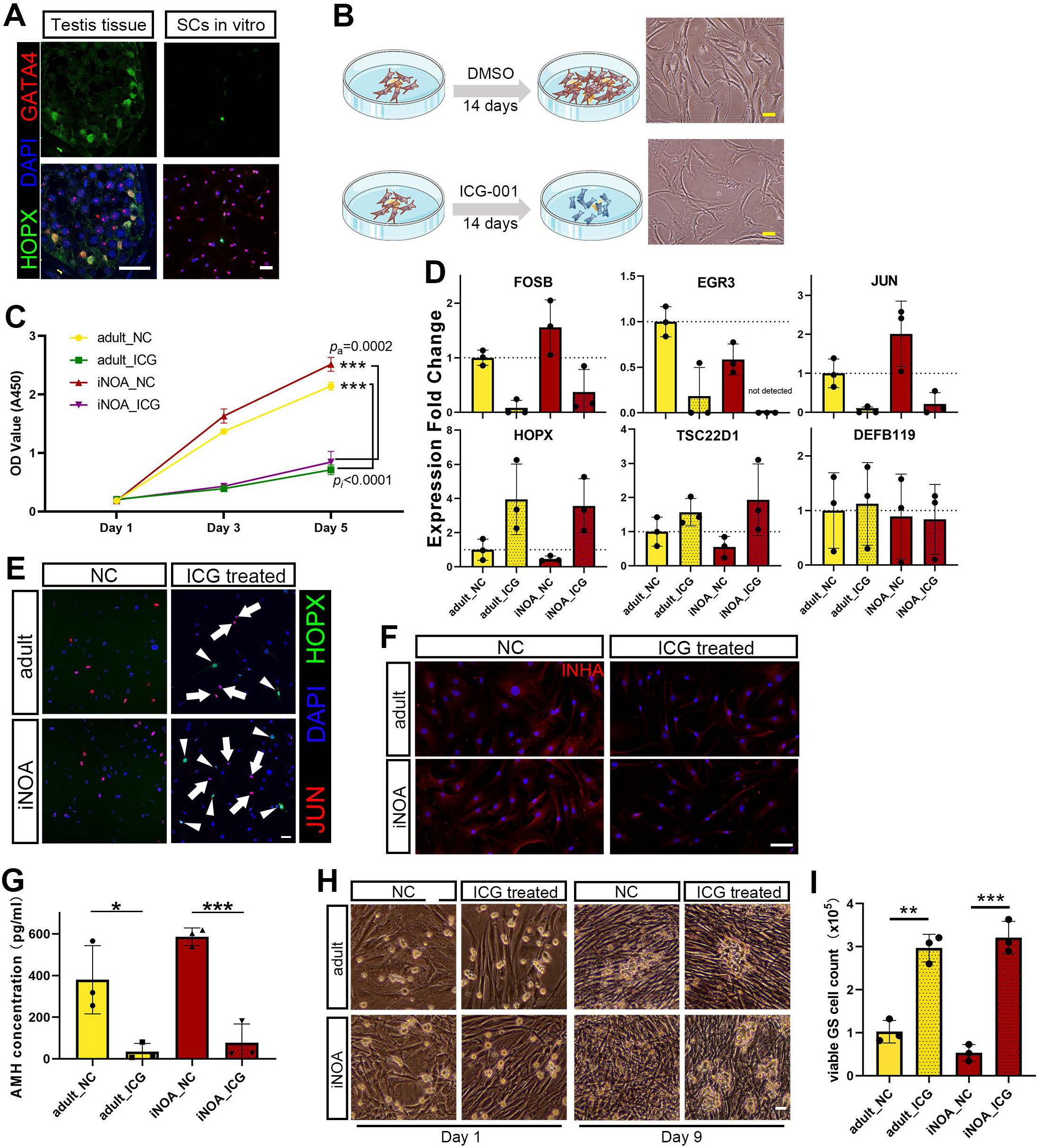
Inhibition of the Wnt pathway stimulated the maturation of Sertoli cells in vitro. (A) Immunofluorescence co-staining of GATA4 (red) and HOPX (green) in normal and pathological testicular paraffin sections and cultured Sertoli cells. The scale bar represents 40 μm. (B) Schematic illustration of the Wnt pathway inhibition experiment. The right panel shows the morphology of cultured Sertoli cells. The scale bar represents 20 μm. (C) Sertoli cells treated with ICG proliferated faster than DMSO-treated controls in vitro, as indicated by CCK-8 assay. (D) qPCR results showing the expression fold change of six stage-specific markers in cultured iNOA and normal adult Sertoli cells with or without ICG treatment. The gene expression levels of normal adult Sertoli cells without ICG treatment were used as the baseline values. (E) Immunofluorescence co-staining of JUN (red) and HOPX (green) in cultured iNOA and normal adult Sertoli cells with or without ICG treatment. The scale bar represents 20 μm. (F) Immunofluorescence of INHA (red) in cultured iNOA and normal adult Sertoli cells with or without ICG treatment. The scale bar represents 20 μm. (G) ELISA results showing the difference in AMH concentration in the supernatant of cultured iNOA and normal adult Sertoli cells with or without ICG treatment. \**P* < 0.05; \*\*\**P* < 0.001. (H, I) The morphology of cultured GS cell colonies when normal adult or iNOA Sertoli cells with or without ICG treatment are used as feeder cells. Images were taken with a 20× magnification microscope.

We next assessed the effects of ICG treatment on the mRNA levels of stage-specific genes by qPCR. We found that Stage_a-specific genes, such as *FOSB, EGR3*, and *JUN*, were downregulated after ICG treatment, while the mRNA levels of Stage_c-specific genes, such as *HOPX* and *TSC22D1*, were significantly higher than in control cells (Figure 6D). However, the expression of *DEFB119*, a highly DEG in Stage_c, was not changed after ICG treatment (Figure 6D), indicating that this gene may be regulated by other stimuli during Sertoli cell development. We further checked the expression changes at the protein level. IHC staining results showed that the proportion of HOPX-positive cells in the ICG-treated group was significantly higher than in the control group. However, the proportion of JUN shown an opposite trend and did not show any co-staining with HOPX (Figure 6E). In addition, the expression levels of INHA and AMH, two ligands of the TGF-β signaling pathway, were low in Stage_c cells, as evidenced by IHC staining and ELISA (Figure 6F-G).

To determine whether the maturity of Sertoli cells affected their support function for germ cells, we used iNOA and normal adult Sertoli cells with or without IGC treatment as feeder cells, and then seeded GS cells, which is a mouse germ cell line mainly composed of SSCs and type A spermatogonia (Zhou et al., 2015). From the 5th day, we observed more and larger GS cell colonies in the ICG-treated group than in the control group. There was no difference in cell colony size between GS cells fed by ICG-treated iNOA cells and ICG-treated normal adult Sertoli cells, but in the GS cells fed by non-treated Sertoli cells, cell colonies fed by normal adult Sertoli cells were observably larger than in the iNOA group (Figure 6H). All GS cells were collected and counted on the 9th day. The counts of viable GS cells in the ICG-treated groups (2.97×105 in the normal adult Sertoli group and 3.21×105 in the iNOA Sertoli group) were also significantly higher than control groups (1.02×105 in the normal adult Sertoli group and 0.53×105 in the iNOA Sertoli group) (Figure 6I), indicating inhibition of the Wnt pathway could alleviate the maturation disorder of iNOA Sertoli cells and improve the supporting function of both iNOA and normal Sertoli cells in vitro.

## Discussion

The role of the testicular somatic microenvironment for germ cells is like the role of soil for sprouting seeds, providing them with various necessary nutrients. Although an increasing body of evidence suggests that the human testicular cells are highly heterogeneous, our understanding of testicular homeostasis and disorders remains hampered by our understanding of cell type complexity and cell-to-cell interactions, especially for somatic cell development. No effective treatments exist for NOA, especially iNOA. This is probably because most NOA patients are not diagnosed until childbearing age, when spermatogenesis has already been irreversibly damaged, and no or few germ cells are left. However, the other testicular cell types, with distinct physiological and pathological characteristics at the molecular level, are also completely absent. In the present study, we compared the developing testes from NOA patients with those of healthy subjects from a new perspective at an unparalleled resolution. Our study is a big step towards the understanding of the testicular microenvironment and male infertility-related diseases.

Most human testicular cell populations functionally develop after birth (Sohni et al., 2019). However, little is known about the development of human testicular cell lineages from newborns to puberty. Our first important finding is that we shed light on the cellular processes underlying the development of the testicular microenvironment by taking developmental time as the horizontal axis. Sertoli cells serve as the scaffold in the microenvironment. Previous studies suggested that Sertoli cells go through only two developmental stages, i.e., from immature to mature (Meroni et al., 2019; Simorangkir et al., 2012). Sertoli cells remain immature until the peak of testosterone production during puberty (Meroni et al., 2019; Tan et al., 2005). Our data, however, show that dramatic changes begin in early childhood, and we found that Sertoli cells go through three independent and consecutive developmental stages (Stage_a, Stage_b, and Stage_c). In infancy and early childhood, Sertoli cells of Stage_a are predominant and proliferate vigorously, with characteristics similar to those of stem cells. This may be due to various factors secreted by early Leydig cells, such as IGF (Cannarella et al., 2019). Rapid amplification helps to increase the volume of seminiferous tubules. Before puberty, proliferation of many Sertoli cells is significantly reduced, and they enter Stage_b. At this stage, Sertoli cells express many genes to prepare for structural maturation. From a metabolic perspective, we observed that once a cell enters Stage_b, its preferred metabolic pathway changes from oxidative phosphorylation to glycolysis. This is consistent with the previous observation that lactic acid (produced by glycolysis) provides energy for germ cell development (Ni et al., 2019). It is worth noting that all three stages of Sertoli cells exist and form a steady state in adult testes, maintaining the basic structure of seminiferous tubules and also guaranteeing homeostasis of the microenvironment for spermatogenesis for decades. Having defined the populations in the testicular microenvironment, we also revealed an unbiased ligand–receptor interaction network among different cell types, providing a more holistic overview. Focusing on Sertoli cells, we investigated the processes underlying the maturation of the microenvironment by analyzing single-cell transcriptomes from humans of different ages.

We systematically compared the construction of the microenvironment in healthy subjects with that in NOA patients. We found that the most common defects observed in NOA patients are dysfunctional Sertoli cells. Other important findings include the main features of Sertoli cell defects in patients with different types of NOA. Both KS and AZFa are caused by a sex chromosome disorder. Because of X-inactivation (Carrel and Willard, 2005), we observed that the total levels of X chromosomal genes in each type of testicular cell in KS are less than twice that in healthy adults; on the other hand, we found Sertoli cells express the highest proportion of X-inactivated genes, indicating Sertoli cells are more susceptible to dysregulation due to an extra X chromosome than other testicular cells. In AZFa_Del patients, although the loss of *DDX3Y* has been reported to cause dysfunction in germ cells, we found *USP9Y*, another AZFa region gene, was specifically expressed in Sertoli cells, indicating the pathological change of AZFa_Del Sertoli cells was not only caused by the loss of germ cells, but also by AZF region deletion. In addition, both KS and AZFa_Del Sertoli cells abnormally expressed high levels of *MIF*, a gene encoding a cytokine that stimulates the immune response and induces apoptosis in immune as well as non-immune cells (Harris et al., 2019). These findings indicate that in NOA patients with genetic disorders in sex chromosomes, Sertoli cells proceed to the mature stage, but a chronic immune response may induce cell death.

In patients with iNOA, we found that Sertoli cells were observably immature and heterogeneous. In addition, over half of the DEGs in cluster 1 overlapped with Stage_a, suggesting that these cells physiologically closely resemble healthy immature Sertoli cells. In cluster 2, we observed an upregulation of genes involved in (i) the immune response pathway and (ii) some Stage_b-specific features, such as structural morphogenesis and the response to steroid hormone. It should be noted that the germ cells themselves also affect the cells in the microenvironment. Could the immaturity of Sertoli cells in iNOA patients simply be caused by the loss of germ cells? We think the answer is negative, for the following two reasons. First, Sertoli cells in AZFa_Del patients without germ cells could nevertheless enter Stage_c. Second, in co-culture experiments with germ cells, we are not able to induce the maturation of patients’ Sertoli cells. Restoring the maturity of the microenvironment may activate local spermatogenic foci. This prompted us to understand the maturation process of Sertoli cells and to screen for diagnostic biomarkers and regulators that could potentially serve as therapeutic targets for iNOA treatment.

Surprisingly, Sertoli cell transcriptomes of patients with iNOA are highly similar, although the possible causes are unknown. Even if the initial causative factors are unclear, finding pathways to improve the microenvironment may still be helpful for disease treatment. We identified some novel candidate pathways and regulators which may play a crucial role during Sertoli cell maturation. The activating signals are mainly stem cell maintenance-related signals, such as the Wnt/β-catenin, ephrin receptor, and integrin pathways. This is consistent with the immature phenotype of Sertoli cells. While the Wnt/β-catenin pathway was activated in all patients, we also found that a negative regulator of the Wnt/β-catenin pathway, HOPX, was not expressed in Stage_a Sertoli cells but highly expressed in Stage_c cells. The Wnt/β-catenin pathway plays an important role in the maintenance of adult organs (including the small intestine and the skin), especially for adult stem cell homeostasis. HOPX is considered to promote stem cell maturation by inhibiting Wnt signaling (Jain et al., 2015). By treating cells with an inhibitor that suppresses Wnt signaling, we successfully promoted the partial maturation of some Sertoli cells derived from patients, and these cells could even maintain the proliferation of spermatogonial stem cells. This is also the first step for the treatment of such patients. Until this proves successful, NOA treatment methods remain very limited, and effective treatment targets are still lacking.

However, RNA-seq cannot answer all the details of the entire process; some functional maturation cannot be evaluated by transcriptome analysis. Orthogonal validation approaches, such as IHC in testes of different ages and gene overexpression/knockdown experiments in animal models, are required. Considering most changes happen in puberty, more intensive sampling around puberty may provide more detailed information about the developmental processes and transitions. Likewise, future studies with larger sample sizes of KS, AZF_Del, and iNOA patients at different ages, especially around puberty, could strongly improve our understanding of the causes underlying spermatogenic dysfunction in these diseases.

In summary, our study not only revealed the major cell types and the processes underlying their maturation processes in normal human testes, but also for the first time clearly compared the maturation disorders of testicular cells in patients with the main types of NOA. These comparisons shed light on the pathogenesis of different types of NOA and pave the way for the development of diagnostic and treatment methods for NOA.

## Methods

### CONTACT INFORMATION FOR REAGENT AND RESOURCE SHARING

Requests for further information (gene expression matrix) and for resources and reagents should be directed to and will be fulfilled by the Lead Contact, Zheng Li.

### EXPERIMENTAL MODEL AND SUBJECT DETAILS

The experiments performed in this study were approved by the Ethics Committee of Shanghai General Hospital (License No. 2016KY196). For human testis samples, all participants (and their legal guardian if aged <18 years) signed their consent after being fully informed of the goal and characteristics of our study. Fresh testicular tissues were obtained from 5 male donors who underwent testicular biopsy or partial excision for the following indications: contralateral testis to testicular torsion (*n* = 1), benign testicle mass (*n* = 3), or contralateral testis to cryptorchidism (*n* = 1), and an additional 5 OA and 7 NOA samples were obtained from the abandoned tissues after testicular sperm extraction operation. Ten normal samples (5 underage and 5 OA donors) all had normal karyotypes, genotypes, sex hormone levels, and morphology of seminiferous tubules according to their age. The KS and AZFa microdeletion donors were diagnosed by spectral karyotyping and RT-PCR. The qPCR examination before hospitalization showed that in this sample, sY84 and sY86 were completely deleted. Among all donors, other abnormal genotypes related to spermatogenic disorders were excluded by whole-exome sequencing.

### Histological examination

Fresh testicular tissues from donors were fixed in 4% paraformaldehyde for 12–24 hours at 4°C, embedded in paraffin, and sectioned. Before staining, tissue sections were dewaxed in xylene, rehydrated using a gradient series of ethanol solutions, and washed in distilled water. Then the sections were stained with PAS/hematoxylin and dehydrated using increasing concentrations of ethanol and xylene. Sections were allowed to dry before applying neutral resin to the coverslips. The staining images were captured with a Nikon Eclipse Ti-S fluorescence microscope (Nikon).

### Immunohistochemical staining

The sections were rehydrated by the same method as for PAS/hematoxylin staining. After rehydration, sections were processed for antigen retrieval with 10% sodium citrate at 105°C for 10 min. Tissues were blocked with 5% normal donkey serum and incubated with appropriate primary antibodies at 4°C overnight. Sections were further incubated with secondary antibody for 2 h at room temperature. Nuclei were labeled with DAPI by incubating tissue sections for 15 min. Images were captured with an OLYMPUS IX83 confocal microscope.

### Isolation of single testicular cells

Testicular cells were isolated from human and mouse testicular tissues by enzymatic digestion according to a previously described method. In brief, testicular tissues were enzymatically digested with 4 mg/ml collagenase type IV, 2.5 mg/ml hyaluronidase, and 1 mg/ml trypsin at 37°C for 20 min. Given the risk of damaging certain types of big cells (such as Sertoli cells), disruption of the samples was avoided during this process. Subsequently, the cell suspension was filtered through a 40-mm nylon mesh, and the cells were sorted by MACS with a Dead Cell Removal Kit (Miltenyi Biotec) to remove dead cells. The cells were re-suspended in 0.05% BSA/PBS buffer before 10× Genomics library preparation.

### Single-cell RNA-seq library preparation

Cell suspensions were loaded on a Chromium Single Cell Controller instrument (10× Genomics, Pleasanton, CA, USA) to generate single-cell gel beads in emulsions (GEMs). Single-cell RNA-seq libraries were prepared using the Chromium Single Cell 3′ Library & Gel Bead Kit (P/N 120237, 10× Genomics) according to the manufacturer’s instructions. Briefly, suspensions containing about 8000 cells per sample were MIX with RT-PCR reaction before being added to a chromium chip already loaded with barcoded beads and partitioning oil. The chromium chip was then placed in a Chromium Single Cell Controller instrument. GEM-RT-PCR was performed in a C1000 Touch Thermal cycler with 96-Deep Well Reaction Module (Bio-Rad; CT022510) to produce barcode cDNA using the following program: 53°C for 45 min; 85°C for 5 min; maintain at 4°C. Barcoded cDNA was isolated from the partitioning oil and then amplified by PCR. Sequencing libraries were generated from amplified cDNA using a 10× chromium kit including reagents for fragmentation, sequencing adaptor ligation, and sample index PCR. Final libraries were sequenced on an Illumina Novaseq 6000.

### Mapping, sample quality control, and integration

Cell Ranger software (version 2.2.0), provided by 10× Genomics, was used to demultiplex cellular barcodes, map reads to the genome and transcriptome using the STAR aligner, and downsample reads as required to generate normalized aggregate data across samples, producing a matrix of gene counts versus cells. After mapping, the filtered count matrices of each sample were tagged with a special library batch ID, and a *Seurat* object was created using the *Seurat* package in R. Cells were further filtered according to the following threshold parameters: the total number of expressed genes, 500 to 9000; total UMI count, between −∞and 35,000; and proportion of mitochondrial genes expressed, <40%. Normalization was performed according to the package manual (https://satijalab.org/seurat/v3.1/pbmc3k_tutorial.html). Because cell capturing with LZ013/LZ014/LZ015 and LZ017/LZ018/LZ019 was performed with a BD Rhapsody system with three samples in one batch, these six samples could be regarded as two independent batches. Other samples captured with 10× Genomics were considered as 11 independent batches. Batch correction was performed using the *IntegrateData* function in the *Seurat* package.

### Cell identification and clustering analysis

The merged *Seurat* objects were scaled and analyzed by principal component analysis (PCA). Then the first 20 principal components (PCs) were used to construct a KNN graph and refine the edge weights between any two cells. Based on all these cells’ local neighborhoods, the *FindClusters* function with the resolution parameter set as 0.1 was used to cluster the cells. In total 16 clusters were identified, and these clusters were renamed by accepted marker genes. The first 20 PCs were also used to perform non-linear dimensional reduction by UMAP, and the dimension reduction plots were given as output (Figure 1C). For further analysis, we isolated Sertoli cells and performed these two steps again; we obtained three clusters.

### Identification of differentially expressed genes

The *Seurat FindAllMarkers* function (test.use = ‘‘wilcox’’) is based on the normalized UMI count to identify unique cluster-specific marker genes. Only the genes that are detected in at least 10% of the cells were tested, and the average log2(fold change) threshold was set as 1 in the analysis of eight testicular cell types and as 0.25 in the subgroup analysis of Sertoli cells. GO analysis was performed using WebGestalt 2019 (http://www.webgestalt.org/#).

### Cell cycle analysis

The cell cycle was analyzed by the function *CellCycleScoring* in the *Seurat* package. Briefly, 43 S phase and 54 G2/M phase genes were used to calculate a score for each cell, and the cell cycle stages of these cells were estimated based on these scores.

### Cell trajectory analysis

Single-cell pseudotime trajectories were constructed with the Monocle 2 package (version 2.8.0) according to the operation manual (http://cole-trapnell-lab.github.io/monocle-release/docs_mobile/). Briefly, the UMI count matrices of the Sertoli cells and Leydig&PTM cells were input as the expr_matrix, and meta.data was input as the sample_sheet. Then, 11521 ordering genes (DEGs among clusters) were chosen to define a cell’s progress. In this step, ordering genes that were expressed in less than 10 cells and had a *P*-value bigger than 0.001 were excluded; batch effects were removed by setting the *residualmodelFormulaStr* parameter. DDRTree was used to reduce the space down to one with two dimensions, and all cells were ordered with the *orderCells* function.

### Cellular energy metabolism analysis

Genes associated with glycolysis, oxidative phosphorylation, or triglyceride metabolism were obtained from the AmiGO 2 database (http://amigo.geneontology.org/). These genes were input into *Seurat*, and the proportions of the genes were calculated by the *PercentageFeatureSet* function.

### Transcription factor network construction

A total of 1,469 human transcription factors in AnimalTFDB and a gene expression matrix were taken as input for the ARACNe-AP software. The results of ARACNe-AP were input into the *ssmarina* (version 1.01) R package, which further calculated the marina objects containing the normalized enrichment scores, *P*-values, and a specific set of regulators.

### Analysis of inactivated X chromosomal genes in the AZFa-Del sample

A list of inactivated X chromosomal genes was obtained from Carrel’s study(Carrel and Willard, 2005). Active and inactive X chromosomal genes were input into the *Seurat* object, and their proportions were calculated by the *PercentageFeatureSet* function.

### Cell proliferation assays

Cell proliferation was analyzed by CCK-8 assay. Sertoli cells were seeded in a 96-well microplate containing 200 μl culture medium per well. After cell attachment, the cells were starved in serum-free medium for 16 h. Then, the medium was replaced with culture medium with 10 μM ICG-001 (ICG treatment) or DMSO (control). After 1 to 5 days of culturing, CCK-8 medium (Dojindo) was added to the cells. The optical density (OD) for each well was measured at 450 nm using a microplate reader (Bio-Rad Model 550).

### GS cell supporting assays

Sertoli cells form OA or iNOA patients were first treated with or without 10 μM ICG-001 in DF12 medium for 14 days. Then 10^5^ Sertoli cells from each group were re-seeded in 6-well microplates as the feeder cells. After 24 h, 10^5^ GS cells were seeded in each well, and the medium was replaced with GS culture medium. After 9 days, GS cells were digested by 0.25% pancreatin for 2 min, and the GS cells were gently aspirated. Digested GS cells were cultured for another 1 h to remove potential Sertoli cells, and stained with Trypan blue before counting.

### AMH secretory ability assay

The AMH concentration in Sertoli cell supernatant was measured with the Human Anti-Mullerian Hormone (AMH) ELISA Kit (JYMBIO) according to the manufacturer’s instructions.

## Acknowledgments

We are greatful to Dr. Lingsong Li and Dr. Yumiko Saga for reading the manuscript. This work was support by grants from National Key R&D Plan of China (2017YFC1002003 and 2018YFA0107702); National Natural Science Foundation of China (81671512, 81701524 and 81701428); the ShanghaiTech University start-up fund of Zhi Zhou; Project funded by China Postdoctoral Science Foundation 2019M661524. We wish to thank the Shanghai OE Biotech CO. LTD and Sinotech Genomics CO. LTD. Shanghai for their support of RNA-seq library preparation.

## Author Contributions

Z.L., Z.Z., C.W. and J.S. conceived the project. L.Z., C.Y., X.X., T.J., C.Y., J.Z., J.L., N.L., Z.D., X.L., J.F. and N.L., performed the experiment. P.L., Z.Z., R.T. and H.C., collected the samples. L.Z. conducted the bioinformatics analyses. L.Z., C.Y. and T.J. wrote the manuscript with help from all of the authors.

