## Supplemental Figure 1-6 for "Single-cell atlas of human developing and azoospermia patients’ testicles reveals the roadmap and defects in somatic microenvironment"

A

| Sample ID | Group | Tech of single-cell capture | Cell Number (After QC: 88723 ) |
|---|---|---|---|
| LZ011 | 2 years | 10X Genomics | 7896 |
| LZ009 | 5 years | 10X Genomics | 4239 |
| LZ005 | 8 years | 10X Genomics | 7515 |
| LZ008 | 11 years | 10X Genomics | 5253 |
| LZ016 | 17 years | 10X Genomics | 5007 |
| LZ003/LZ007 | 23/28 years (OA) | 10X Genomics | 6342/8840 |
| LZ013/LZ014/LZ015 | 25/28/31 years (OA) | BD Rhapsody | 3197/3467/3530 |
| LZ002 | 31 years (AZFa Microdeletions) | 10X Genomics | 5203 |
| LZ004/LZ010/LZ012 | 26/27/29 years (Klinefelter's Syndrome) | 10X Genomics | 2743/2740/2244 |
| LZ017/LZ018/LZ019 | 32/26/31 years (NOA) | BD Rhapsody | 7402/6526/6579 |

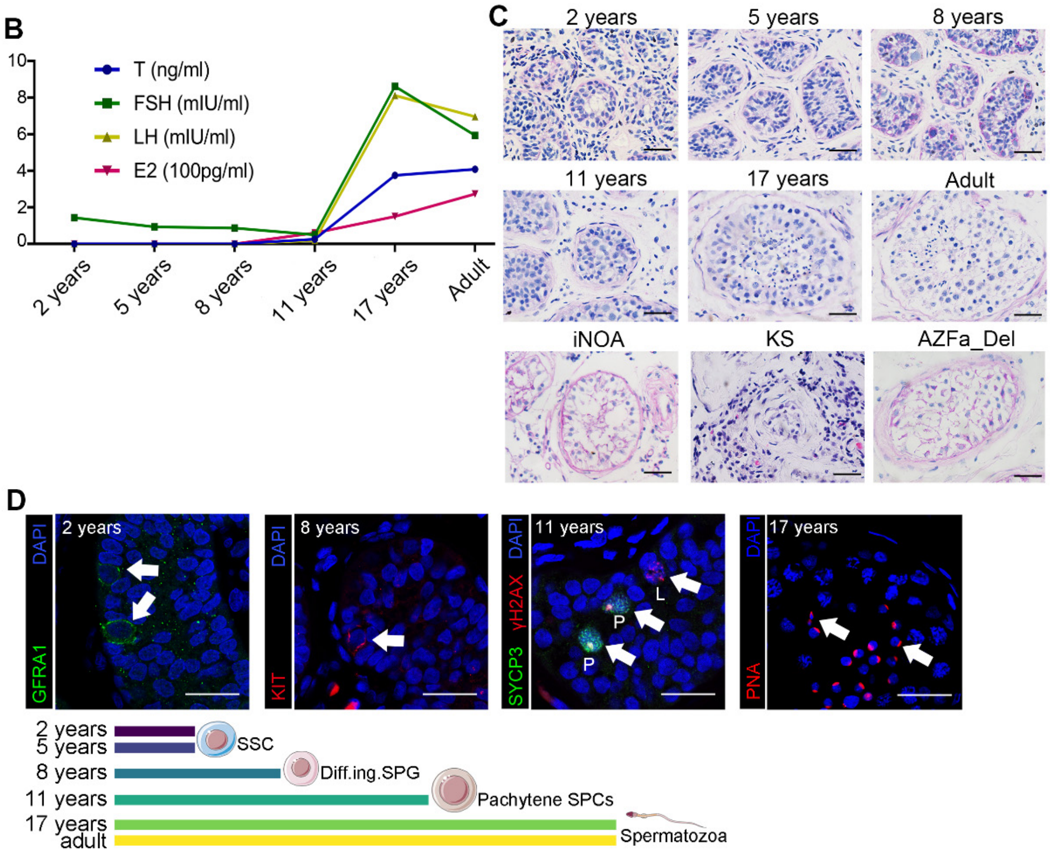


### Figure S1. Clinical and histological information of enrolled samples.

(A) Clinical information of enrolled samples in this study.

(B) Sex hormone levels at each age. The sex hormone levels of adults are presented as the mean of 5 adult samples. T, testosterone; FSH, follicle-stimulating hormone; LH, luteinizing hormone; E2, estradiol.

(C) Periodic acid–Schiff (PAS) staining of each sample enrolled in this study illustrated the normal maturation and pathological changes in histological morphology with age.

(D) Immunofluorescence staining for GFRA1 (SSC marker), KIT (differentiating SPG marker), and PNA (spermatid marker) and double staining of SYCP3 and γH2AX (SPC markers) in testicular paraffin sections of each age, illustrating the spermatogenic maturity in each sample. The scale bar represents 50 μm.

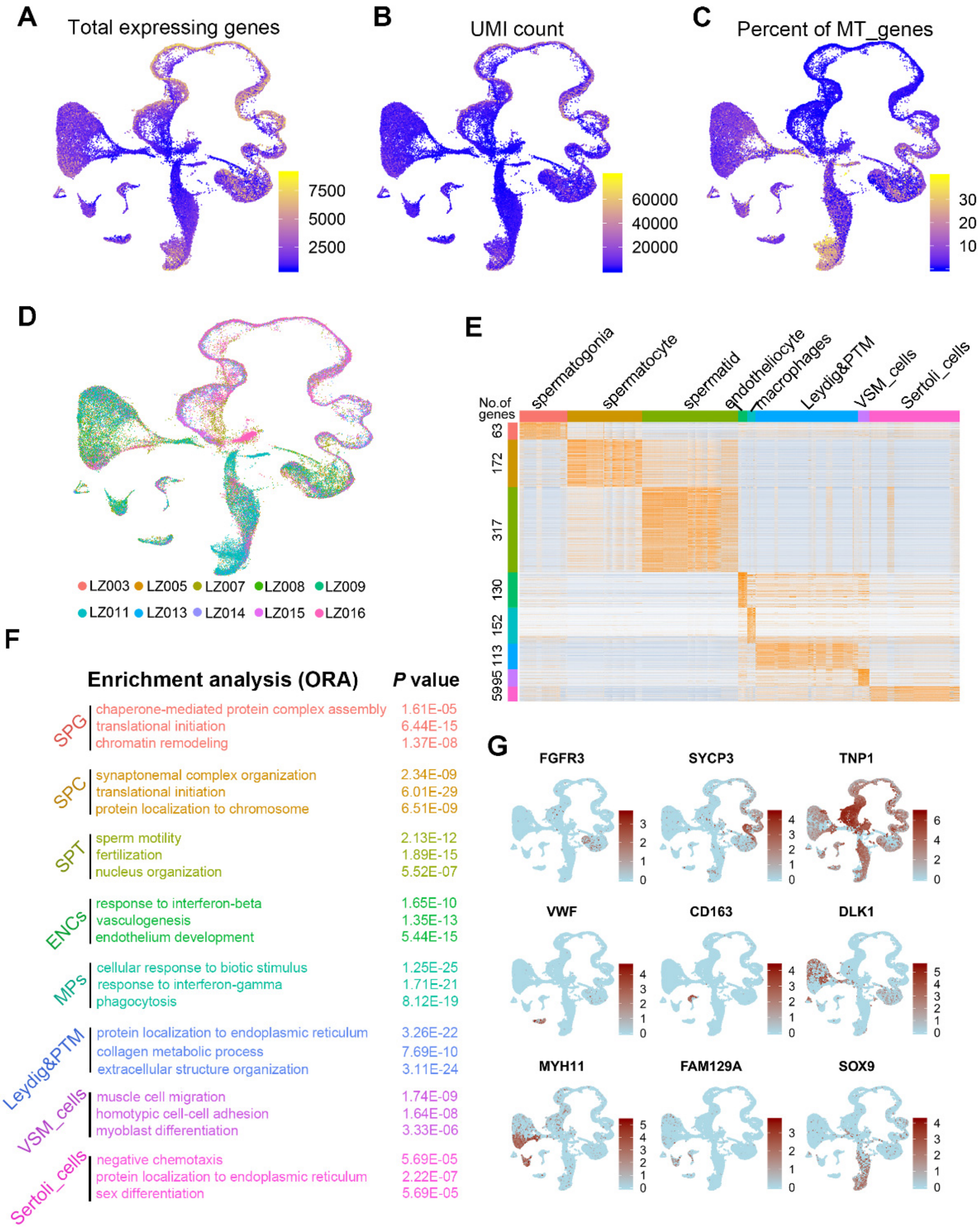


Figure S2. Quality control of single-cell RNA-seq datasets and identification of nine clusters of normal developing testicular cells.

(A–C) Single-cell RNA-seq quality information, including (A) total count of expressed genes, (B) UMI counts, and (C) percentage of mitochondrial gene expression, is projected on the UMAP plot.

(D) UMAP plots of all 10 healthy human testicular cell samples. Each sample is labeled with a different color.

(E) Heatmap showing the DEGs of each cluster. DEG counts are shown on the left of the color bar of the cell type annotation.

(F) Enriched GO terms and *P*-values of each cell cluster are indicated by eight distinct colors.

(G) Expression patterns of the following markers for each cluster are projected on the UMAP plot: FGFR3 (spermatogonia), SYCP3 (spermatocytes), TNP1 (spermatids), VWF (endotheliocytes), CD63 (macrophages), DLK and MYH11 (MIX cells), FAM129A (vascular smooth muscle cells), and SOX9 (Sertoli cells). A gradient of light blue to dark red indicates low to high expression levels.

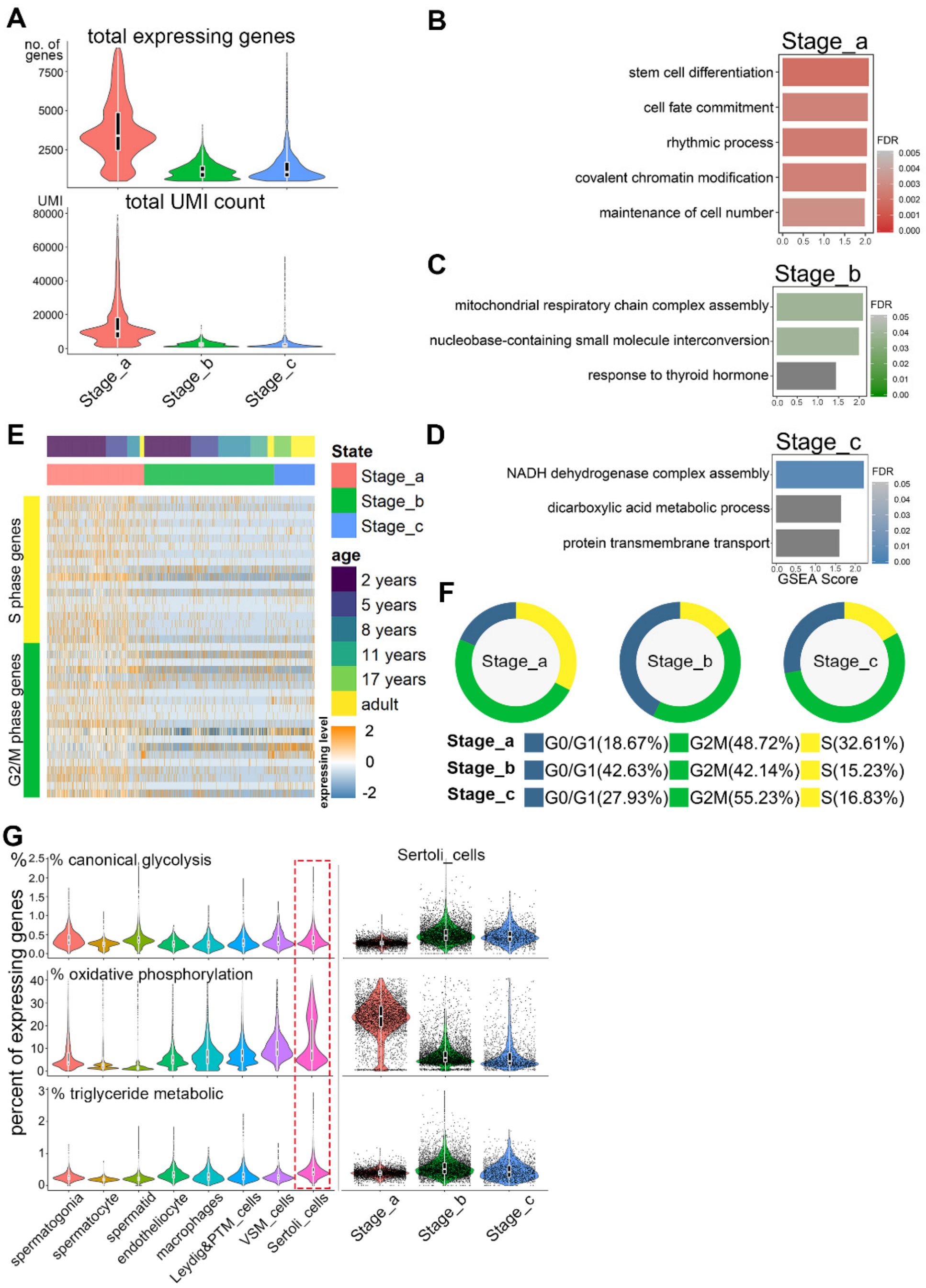


Figure S3. Characteristics related to the cell cycle, energy metabolism, and biological processes of each stage during Sertoli cell development.

(A) Violin plot of overall transcriptional levels of Sertoli cells at each stage.

(B–D) Pathways and biological process terms of GSEA are shown as barplot. The GSEA score is presented on the *x*-axis, and a gradient of three colors indicates low to high FDR values of the three stages.

(E) Heatmap of cell cycle-specific genes in Sertoli cells at each stage.

(F) Cell cycle score of Sertoli cells at each stage.

(G) Violin plot of the expression levels of energy metabolism-related genes of nine types of testicular cells (left panel) and Sertoli cells at each stage (right panel).

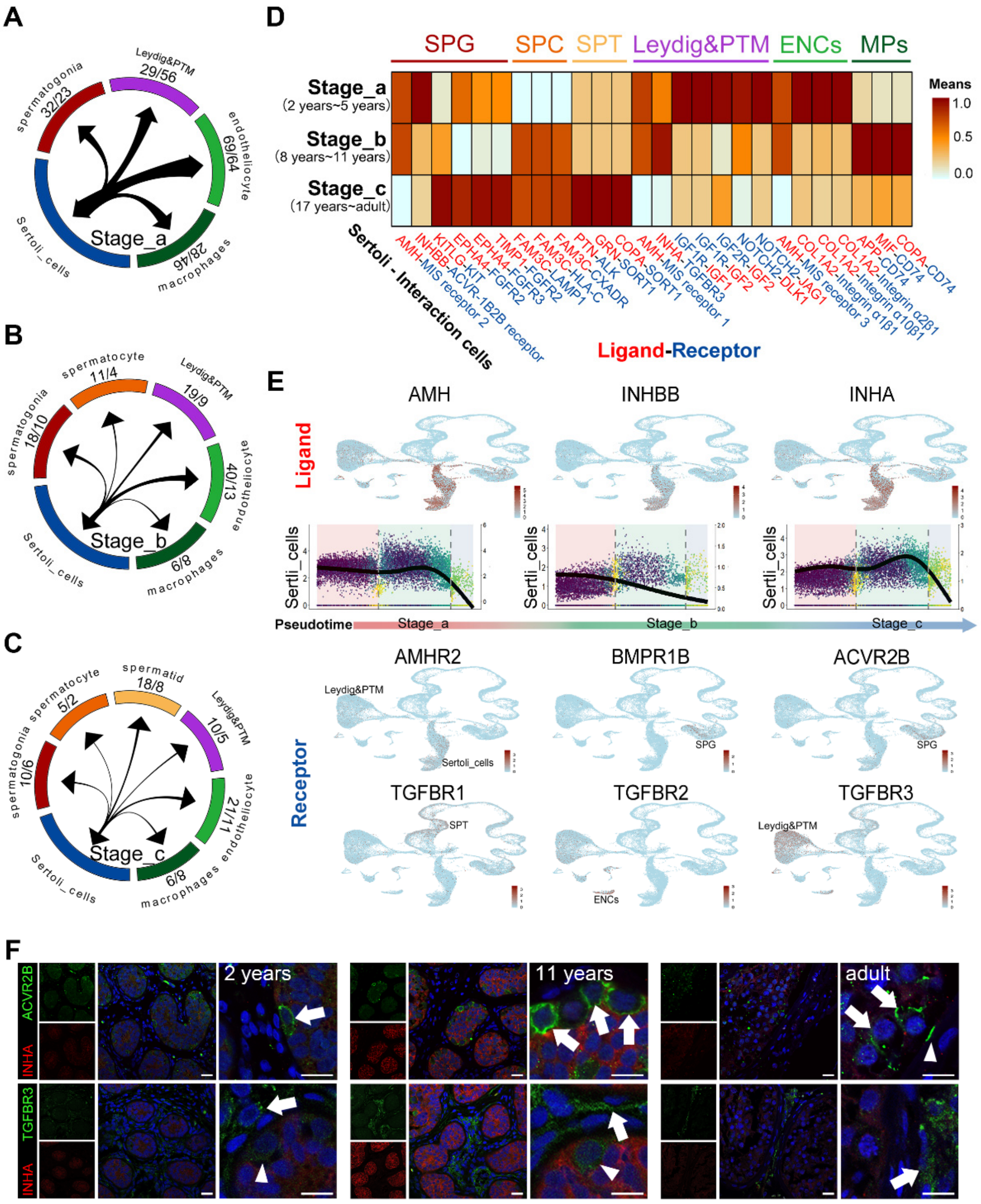


### Figure S4. Dynamic changes of the interactions between Sertoli cells and other testicular cells.

(A–C) Loop graph showing the number of ligands–receptors interactions between Sertoli cells at each stage and other testicular cells. The numbers of ligands/receptors from Sertoli cells are shown under the cell type annotation.

(D) Heatmap showing the matching strength of the interaction between Sertoli cells and other testicular cells.

(E) Analysis of TGF-β signaling (UMAP plot showing the spatial expression pattern of ligands and receptors of the TGF-β signaling pathway, point plot showing the temporal expression patterns of ligands secreted by Sertoli cells).

(F) Immunofluorescence co-staining of INHA (red) with ACVR2B (green, upper panel) and TGFBR3 (green, lower panel) in human testicular paraffin sections at three ages. The scale bar represents 15 μm.

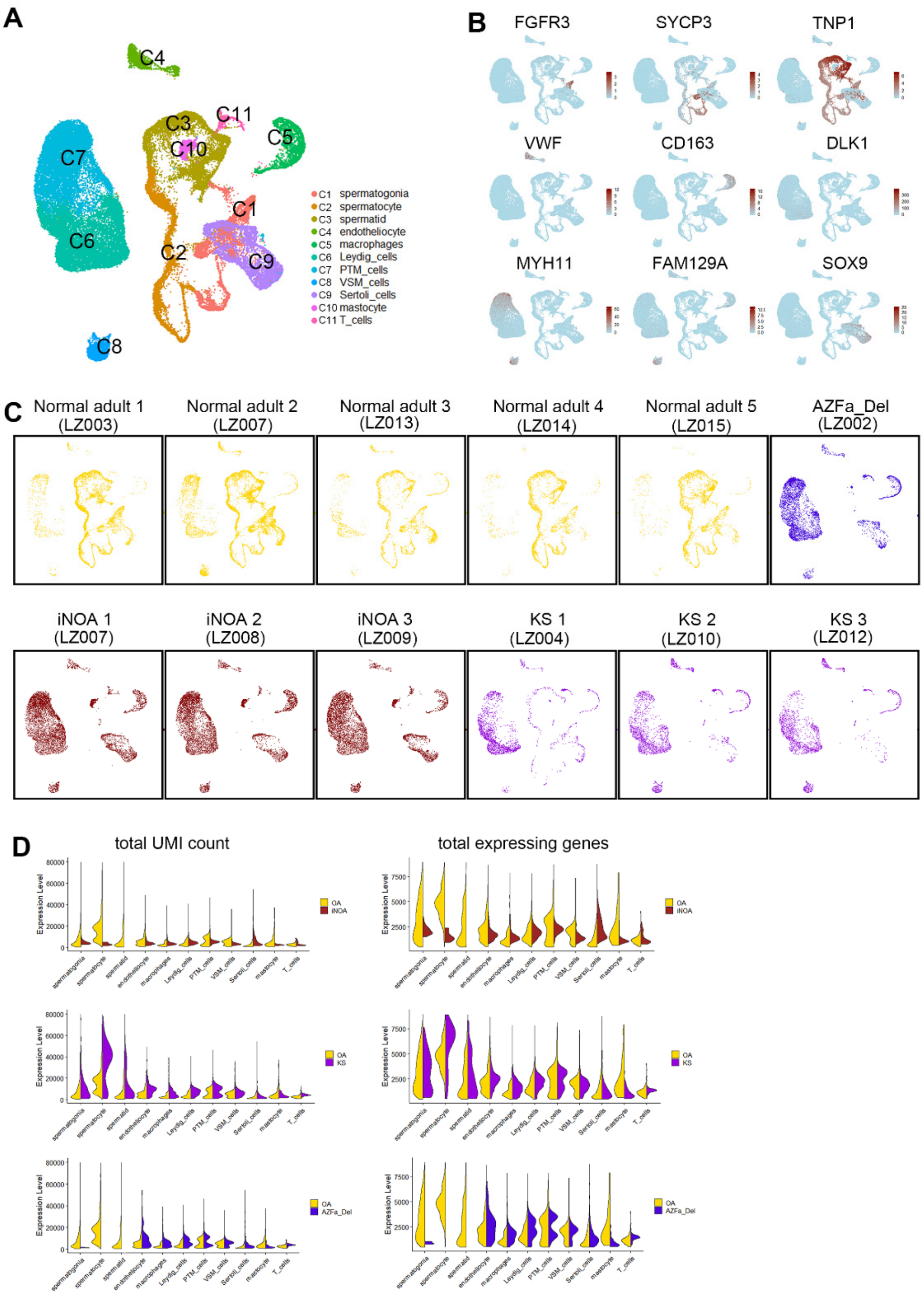


Figure S5. The heterogeneity among different types of NOA and the repeatability with samples of the same type of NOA.

(A) UMAP plots of 10 healthy samples combined with three types of NOA.

(B) Expression patterns of the following markers for each cluster are projected on the UMAP plot: FGFR3 (spermatogonia), SYCP3 (spermatocytes), TNP1 (spermatids), VWF (endotheliocytes), CD63 (macrophages), DLK and MYH11 (MIX cells), FAM129A (vascular smooth muscle cells), and SOX9 (Sertoli cells). A gradient of light blue to dark red indicates low to high expression levels.

(C) UMAP plot of each isolated sample. Healthy and different NOA cells are colored differently.

(D) Violin plot of overall transcriptional level of different cell clusters. Healthy cells and different NOA clusters are split in different columns.

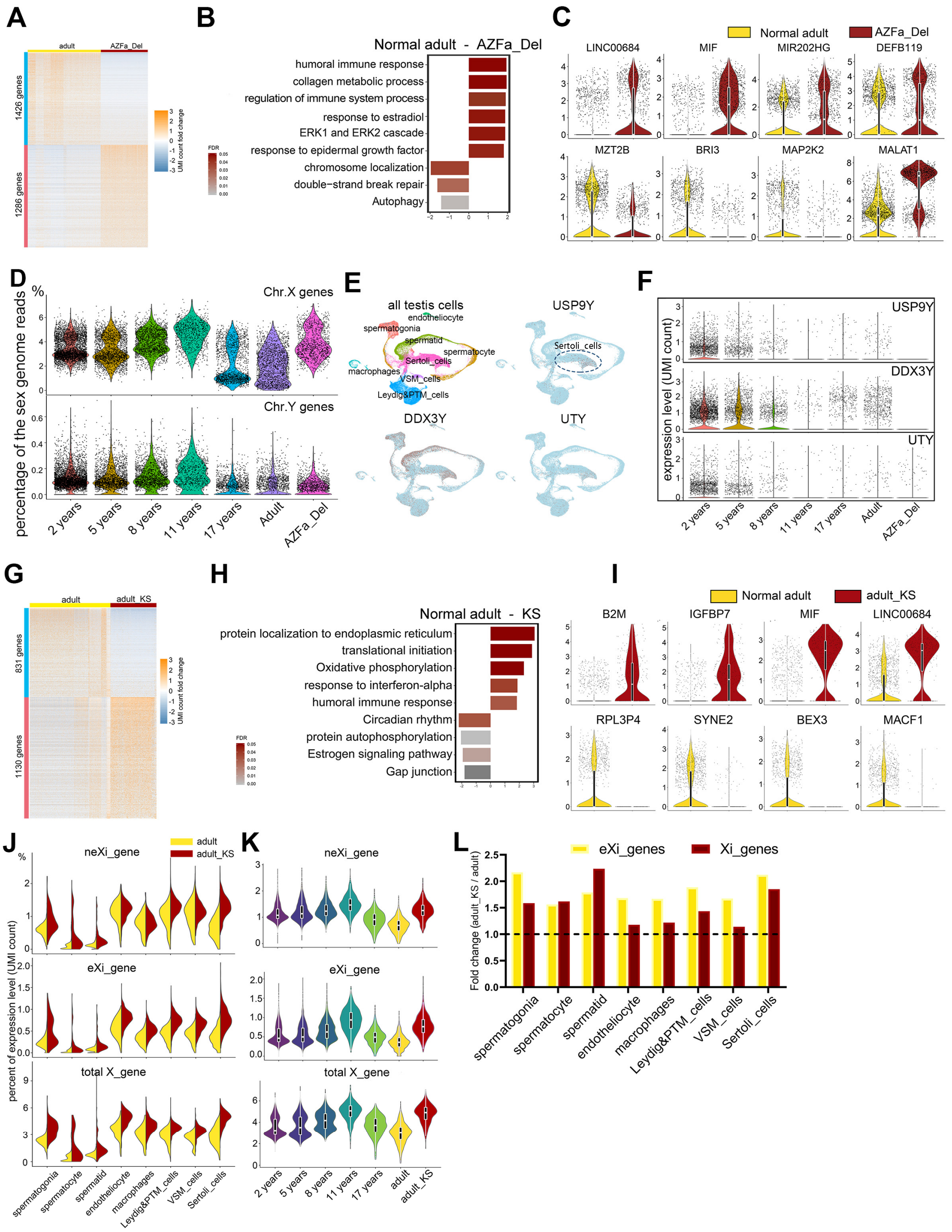


### Figure S6. Abnormal expression patterns in AZFa_Del and KS.

(A) Heatmap of DEGs between healthy adult Sertoli cells and AZFa_Del Sertoli cells.

(B) Violin plot of the expression levels of the top four positive and negative DEGs in AZFa_Del Sertoli cells.

(C) GSEA terms enriched and decreased in AZFa_Del Sertoli cells, shown as barplot. The GSEA score is presented on the *x*-axis, and the gradient of red indicates low to high FDR values.

(D) Violin plot of the expression levels of sex chromosomal genes at in normal Sertoli cells (six different ages) and AZFa_Del Sertoli cells.

(E) UMAP plot showing the expression patterns of three genes in the AZFa region.

(F) Violin plot of the expression levels of three AZFa genes in normal Sertoli cells (six different ages) and AZFa_Del Sertoli cells.

(G) Heatmap of DEGs between normal adult Sertoli cells and KS Sertoli cells.

(H) Violin plot of the expression levels of the top four positive and negative DEGs in KS Sertoli cells.

(I) GSEA terms enriched and decreased in KS Sertoli cells, shown as barplot. The GSEA score is presented on the *x*-axis, and the gradient of red indicates low to high FDR values.

(J, K) Violin plot of the expression level of eXi and neXi genes (J) in eight types of testicular cells and (K) in normal (six different ages) and AZFa_Del Sertoli cells.

(L) Barplot showing the ratio of the expression levels of eXi (yellow) and neXi (dark red) expression levels between healthy adults and KS patients.
